# Multiplex Confounding Factor Correction for Genomic Association Mapping with Squared Sparse Linear Mixed Model

**DOI:** 10.1101/228114

**Authors:** Haohan Wang, Xiang Liu, Yunpeng Xiao, Ming Xu, Eric P. Xing

**Affiliations:** Language Technologies Institute, School of Computer Science Carnegie Mellon University, Pittsburgh, PA, USA; School of Information and Communication Engineering Beijing Univ. of Posts & Telecoms, Beijing, China; Chongqing Engineering laboratory of Internet and Information Security, Chongqing Univ. of Posts & Telecoms, Chongqing, China; Research Institute of Information Technology Tsinghua University, Beijing, China; Machine Learning Department, School of Computer Science Carnegie Mellon University, Pittsburgh, PA, USA

## Abstract

Genome-wide Association Study has presented a promising way to understand the association between human genomes and complex traits. Many simple polymorphic loci have been shown to explain a significant fraction of phenotypic variability. However, challenges remain in the non-triviality of explaining complex traits associated with multifactorial genetic loci, especially considering the confounding factors caused by population structure, family structure, and cryptic relatedness. In this paper, we propose a Squared-LMM (LMM^2^) model, aiming to jointly correct population and genetic confounding factors. We offer two strategies of utilizing LMM^2^ for association mapping: 1) It serves as an extension of univariate LMM, which could effectively correct population structure, but consider each SNP in isolation. 2) It is integrated with the multivariate regression model to discover association relationship between complex traits and multifactorial genetic loci. We refer to this second model as sparse Squared-LMM (sLMM^2^). Further, we extend LMM^2^/sLMM^2^ by raising the power of our squared model to the LMM^*n*^/sLMM^*n*^ model. We demonstrate the practical use of our model with synthetic phenotypic variants generated from genetic loci of Arabidopsis Thaliana. The experiment shows that our method achieves a more accurate and significant prediction on the association relationship between traits and loci. We also evaluate our models on collected phenotypes and genotypes with the number of candidate genes that the models could discover. The results suggest the potential and promising usage of our method in genome-wide association studies.

## I. Introduction

Genome-wide Association Study (GWAS) has been used to relate a number of causal variants and trait variables for a long time and many relations have been revealed, such as the genetic architecture of global level traits in plants [1] and mice [2], also risks for human diseases, like type rheumatoid arthritis [3]. However, it is widely recognized to be challenging for statistical analysis to understand associations because of the difficulty raised by small individual effects and many-to-many gene-trait relationships. Further, confounding relatedness between samples inherently limits the power to unveil weak effects. It is necessary to address these challenges at the same time, modeling combinatorial associations while correcting population and genetic confounding factors, in order to understand the true genetic architecture of complex traits.

Despite the achievement made so far with simple methods that assess the significance of individual loci independently, these independent testing methods do not reach genome-wide mapping power due to the belief of multiple variants contributing to phenotype variation in an additive manner [4]. The challenge of additive effects of multiple SNPs has been widely addressed with multivariate regression of all genome-wide SNPs, with a shrinkage prior or stepwise forward selection [5], Laplace prior [6] and its extension [7], or other complicated modern priors [8], [9], [10].

However, the aforementioned methods easily fall into the trap of another challenge: the population and genetic structure may induce spurious correlations between genotype and phenotype. To address this challenge, Principle Component Analysis that extracts the major axes of population differentiation from genotype data has been explored [11]. Linear mixed model is another tool that provides a more dedicated control of modeling population structure with its random effect component and it has been shown to greatly reduce the impact of population structure [12], [13], [14], [15], [16], [17], [18], [19], [20].

Then a natural following question is about combining the above methods and addressing these two challenges simultaneously. As a result, Segura *et al.* proposed a related multi-locus mixed model approach with the stepwise forward selection [21]. Rakitsch *et al.* proposed a method that bridges the advantages of linear mixed models together with Lasso regression [22]. Additional works proposed to extend previous LMM-Lasso and incorporate more regularizers to further improve the performance in modeling combinatorial associations. [20], [23] These methods grant us the chance to consider complex genetic effects while reducing the impact caused by population structure. However, there might be other cryptic relatedness that cannot be effectively corrected by current implementation of LMM.

In this paper, we propose a new way of constructing the Kinship matrix for a better modeling of cryptic relatedness in addition to population structure and family structure with the introduction of squared LMM (LMM^2^). We further extend our idea to SLMM^2^, LMM^*n*^, and SLMM^*n*^. In the analysis on synthetic data with generated phenotypic variants and real genetic variants from Arabidopsis Thaliana, we show that our model can effectively correct confounders and discover the associated markers.

The contributions of this paper are three-fold:

- We introduce an extension of LMM by raising the power of the kinship matrix, leading to two direct implementations LMM^2^ and sLMM^2^ dependent on the testing procedure. We further extend these two models to even higher power.
- Our proposed models can also be seen as a proof of the concept of the limitation of current realization of kinship matrix as *ZZ^T^*.
- Despite the seemingly simple extension, we offer substantial arguments in both statistical view and genetic view for the necessity of raising the power of kinship matrix. These arguments are further validated with simulations and real data experiments.

## II. Method

### A. Linear Mixed Model (LMM)

For an LMM, suppose we have *m* samples, with phenotypes *y* = (*y*_1_,*y*_2_,…,*y_m_*) and genotypes *X* = (*x*_1_,*x*_2_,…*x_m_*) and for each *i* = 1,2,…, *m*, we have *x_i_* = (*x_i_*,_1_, *x_i_*,_2_,…*x_i,s_*), i.e., each sample has *s* markers to be tested. With linear mixed model, we have:

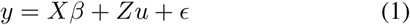

where *β* stands for genetic effect, *u* stands for random effects 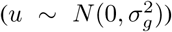, *Z* stands for population structure, and *∈* stands for observation noise 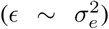. *Z* is not always directly observed from data, but can be conveniently achieved as *Z* = *X*, which is a convention that is widely used in GWAS research (e.g. [14], [24]).

Equivalently, Equation 1 can be formalized as following:

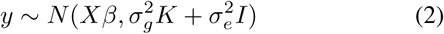

where *K* = *ZZ^T^*. K is also called kinship matrix.

The vanilla LMM assumes 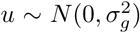 to be identically and independently distributed across *Z*.

### B. Squared Linear Mixed Model (LMM^2^)

This i.i.d assumption of *u* mentioned above might not be general enough. For example, the assumption is justifiable when *Z* encodes the family structure since there are barely inter-family influences that can confound the phenotypes. However, when *Z* encodes pedigree information, there can be correlations of the random effects introduced via the underlying relationship of family tree. This is especially true when we use the genome to denote population structure, i.e when *Z* = *X*, in which case, the existence of LD will make the naive assumption even more problematic.

To account for this, we extend LMM by relaxing the i.i.d assumption of u and propose another assumption that 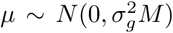, where the covariance matrix *M* describes the correlation induced by confounding factors. Interestingly, *Z^T^Z*, as the covariance matrix, is a reasonable choice for *M*. Therefore, our proposed extension of Equation 2 leads to:

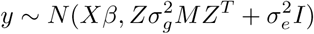

which is:

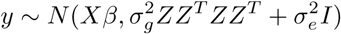

and this leads to an elegant result that our model can be represented as:

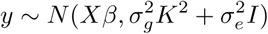

which accounts to squaring the kinship matrix of original LMM. Therefore, we refer to this model as LMM^2^.

Additionally, considering the confounding factors that caused by cryptic relatedness that cannot be explained with the above-accounted relationships like population structure, family structure or genetic correlation, we explore to raise the power of squared LMM further. As an extension of squared LMM, we propose a LMM^*n*^ model that could automatically learn the order *n* of kinship matrix from data with likelihood maximization. Formally, LMM^*n*^ is as following:

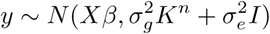

#### 1) Statistical Advantage

In addition to the convenient derivation showed above, another natural statistical advantage introduced via higher order kinship matrix is the ability to distinguish random effects from fix effects. Since *Z* is set as *X* in most cases [14], [24]. By setting *K* = *ZZ^T^*, we will have

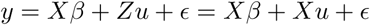

This formulation will lead to the identification problem for the method to differentiate *β* out of *u*.

With higher order of kinship matrix, the method now models the data as:

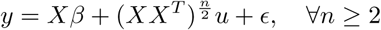

Therefore, the method naturally avoid the identification problem between *β* and *u*.

#### 2) Higher Order Kinship Matrix

In addition to the elegant mathematical derivation of LMM^2^ and extensions to LMM^*n*^, we continue to discuss the intuitive understanding of higher order LMM.

As a simple example, we consider a kinship matrix that describes the pedigree information of individuals as showed in Figure 1(a). Figure 1 (a) shows the direct relationship collected. Each node stands for one individual. The connection between each individual means that the above individual is the parent of the below one. Obviously, kinship matrix here may miss some information as it does not consider any implicit connection. A squared kinship matrix could infer second order relationship, as showed in Figure 1 (b). Now, sibling information and grandparental information are modeled. Figure 1 (c) shows a even higher order kinship matrix, where further related information can be accounted. This example shows the necessity of raising raising the order of kinship matrix as higher order kinship matrix can reveal implicit information.

**Figure 1:**
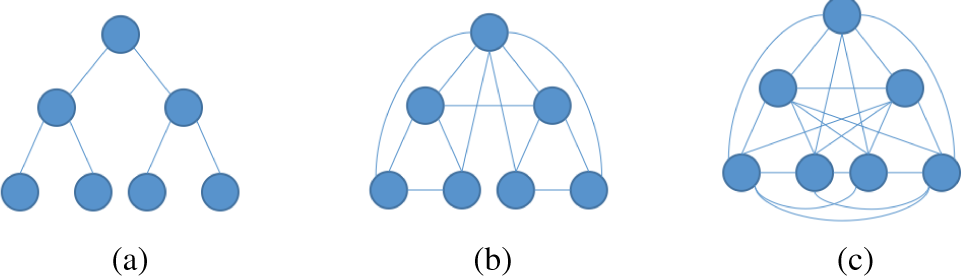
Different order of kinship matrix captures the different information.

This example leaves us with two questions: 1) Whether it is always helpful to raise the order of kinship matrix. 2) Whether the order of kinship matrix is the higher the better.

We proceed to answer these two questions with another example. Assuming we have 20 individuals from three populations as showed in Table I.

**Table I:** A simple example that contains three different populations of individuals, categories are decided according to their different percentage of genome inherited from different continents.

| Group ID | Group 1 | Group 2 | Group 3 |
| --- | --- | --- | --- |
| European | 0.1 | 0.4 | 0.5 |
| American | 0.3 | 0 | 0.5 |
| African | 0.6 | 0.4 | 0 |
| Asian | 0 | 0.2 | 0 |

As a start, we simply represent the population of each individual as a scalar, then *Z* is a 20×1 vector, and the kinship matrix can be calculated as *ZZ^T^*. Figure 2 shows the kinship matrix under different orders and we could see that raising the order of kinship matrix does not gain any improvement.

**Figure 2:**
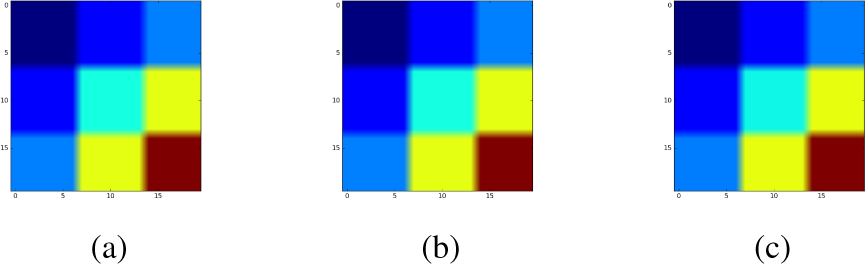
Different order of kinship matrix captures the same information when kinship matrix does not capture the relationship of different groups. (a) first order kinship matrix. (b) second order kinship matrix. (c) kinship matrix when it is converged.

Then, we directly use the percentage to represent the group information, therefore, group information is described with a 20 matrix *Z* and kinship matrix is *ZZ^T^*. Raising the order of kinship matrix can allow the model to consider different relationships between different groups, as showed in Figure 3.

**Figure 3:**
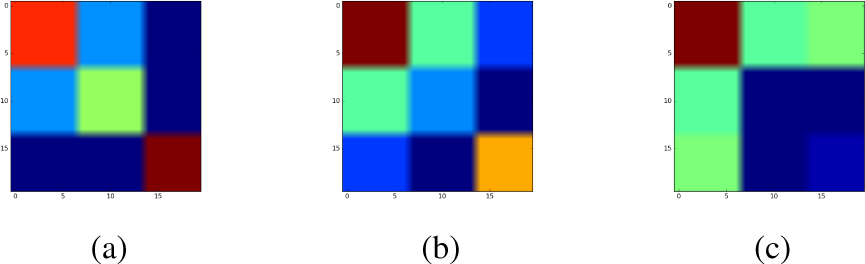
Different order of kinship matrix captures the different information when kinship matrix captures relationship of different groups. (a) first order kinship matrix. (b) second order kinship matrix. (c) kinship matrix when it is converged.

However, when the kinship matrix is converged to a very high order, the relationship is not very well captured since Figure 3(c) cannot distinguish 2^nd^ and 3^rd^ population well. Therefore, raising the order infinitely high until it converges may not be the best practice.

At this point, we have mathematically and intuitively explained the advantages of higher order kinship matrix, which supports the idea of LMM^2^ and LMM^*n*^. We now proceed to introduce the idea of sLMM^2^ and sLMM^*n*^ after we briefly discuss the idea of sLMM.

### C. Sparse Linear Mixed Model (sLMM)

To also account for multifactorial association mapping, recent works proposed a sparse Linear Mixed Model called LMM-Lasso [22], [20], where they added a Laplacian shrinkage prior over the fixed effect *β* and as a result, the majority of genetic effects will be zero.

In detail, instead of traditional likelihood function of LMM, sLMM solves:

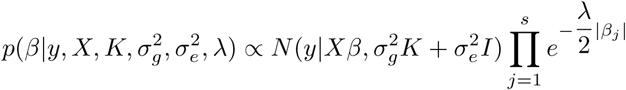

where λ is the sparsity hyperparameter of the Laplace prior.

### D. Squared Sparse Linear Mixed Model (sLMM^2^)

The sparse Squared-LMM (sLMM^2^) is a simple extension of sLMM, with a non-trivial belief that such an extension could effectively correct confounders caused by population structure and genetic structure simultaneously, as extensively discussed in Section II-B2.

Instead of 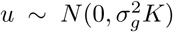 our model believes that the random effect variable 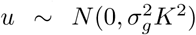 and the model becomes:

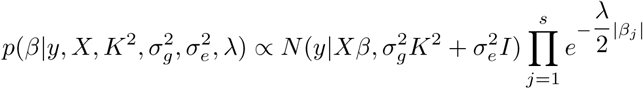

### E. Ordered-N Sparse Linear Mixed Model (sLMM^*n*^)

sLMM^*n*^ extends sLMM^2^ by relaxing the order of covariance component of *u*. Therefore 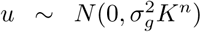. Different from sLMM^2^ where *K*^2^ could be calculated beforehand, the challenge of SLMM^*n*^ lies on how to reliably estimate *n*.

### F. Parameter Estimation

We extend the parameter estimation procedure introduced in [20], where we first correct the confounders with a null model and then apply either independent hypothesis testing or combinatorial sparse regularized multivariate regression afterwards. The estimation of n is conducted via a Grid Search algorithm in the first phase of the algorithm when Brent search is conducted to search for optimal variance parameters.

An important point is that, raising the power of *K* does not significantly increase the computation load because we can utilize the fact that the estimation procedure requires eigendecomposition of *K*, and trivially raise the order of *K* by *K^n^* = *US^*n*^V^T^*, where *K* = *USV^T^* is the eigendecomposition.

## III. Results

We perform experiments to compare the performance of raising the power of kinship matrix. More specifically, we compare the performance among LMM, LMM^2^ and LMM^*n*^, we also compare the difference between sLMM, sLMM^2^ and sLMM^*n*^. By analyzing the results, we reach the conclusion that when single association testing method behaves best and when combinatorial association method will lead to a better result. For LMM^*n*^ and sLMM^*n*^, we set *n* ∈ {3,4,5} in application to avoid the situation when higher order models only behave similar to vanilla case or squared case for a better understanding of necessity of even higher power.

We do not consider other LMM methods in addition to the vanilla LMM because the main contribution of our method is raising the power of the kinship matrix. Other methods, including [17], [19], [18], can always raise the corresponding Kinship matrix to achieve a similar performance. In other words, the novelty of our paper can also be understood as introducing a general framework with the argument that raising the power of kinship matrix can effectively increase the power of confounding correction. However, we do not have to exhaust every LMM implementation to validate this point. Instead, this paper only concerns with the experiments regarding vanilla LMM, sLMM and their corresponding higher order counterparts to prove the concept of higher order kinship matrix.

We compare these models with three different experiments. The first two experiments are performed on semi-empirical synthetic data sets, out of which, the first experiment is to show that different order of models can consider different confounding structure and the second experiment is a set of repeated experiments on a variety of different configurations for semi-empirical synthetic data sets. These repeated experiments are to show that higher order models can consistently perform better than traditional settings as they consider more information.

The third experiment is based on real data set. Following the standard in [22], to handle the non-availability problem of gold standard data set of genotype-phenotype association relationship, we test our results with candidate genes. If the discovered SNP belongs to a candidate gene for the phenotype, this discovery is seen as a true positive discovery.

### A. Arabidopsis thaliana

The Arabidopsis thaliana data set we obtained is a collection of around 1300 plants, each with around 215k SNPs [25]. The latitude and longitude for each of these plants are also available. The candidate genes are collected from [1], we considered eight candidate gene sets, which correspond to 28 phenotypes.

### B. Semi-emperical Synthetic Data Set Experiment One

First, we show that to raise the order of Kinship matrix can gain us the benefit of handling complex confounding structures once they exist.

#### Synthetic Data

To show that, we generate two synthetic data sets based on real SNPs. Without loss of generalizability, we only consider the SNPs of Chromosome 1, which are denoted as *X*. There are about 50k SNPs considered (*p* = 50,000). 100 causal (*N* = 100) SNPs are randomly sampled and fixed effect of causal SNP is uniformly sampled from 0 to 1, namely:

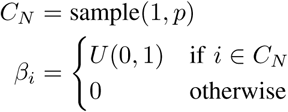

Two synthetic phenotypes are generated as follows:

a. With Population structure: Population structure of plants are collected by clustering their genome sequences. We cluster these plants into 100 groups, each group (Group *k*) is assigned with an effect, namely:

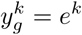

where *K* is the index of clustered group and e is the effect sizes that are sampled from i.i.d, i.e. *e^k^* ∼ *N*(0,1), ∀*k* Then, the phenotype is generated as follows:

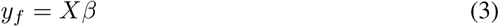

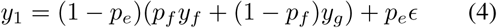

where *p_e_* stands for observation noise effect, *p_f_* stands for the weight of fixed effect, so 1 — *p_f_* can be interpreted as heritability.
b. With Complex Population Structure: In addition to Case (a), we introduce higher order group structure, which simulates the fact when the effect sizes in Case (a) are not i.i.d. but governed by the group structure. Other than clustering *X* into groups in Case (a), we clustered *XX^T^* into groups so simulate complex population structure. Therefore, we have the phenotypes as following:

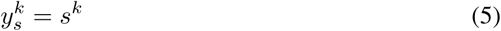

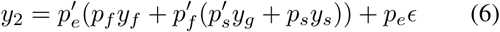

where, same as Case (a), *s^k^* stands for the kth value of a pre-assigned vector of complex population effects and *p_s_* stands for the weight of complex population effect, and 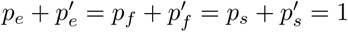 We generate *y*_1_, *y*_2_ by setting *P_e_* = 0.1, *P_f_* = 0.6 and *P_s_* = 0.7 across the data generation process.

#### 1) Experiment Results

The results are showed as a comparison of ROC curve. Here, we truncate the curve to show the differentiating part of curves, as showed in Figure 4.

**Figure 4:**
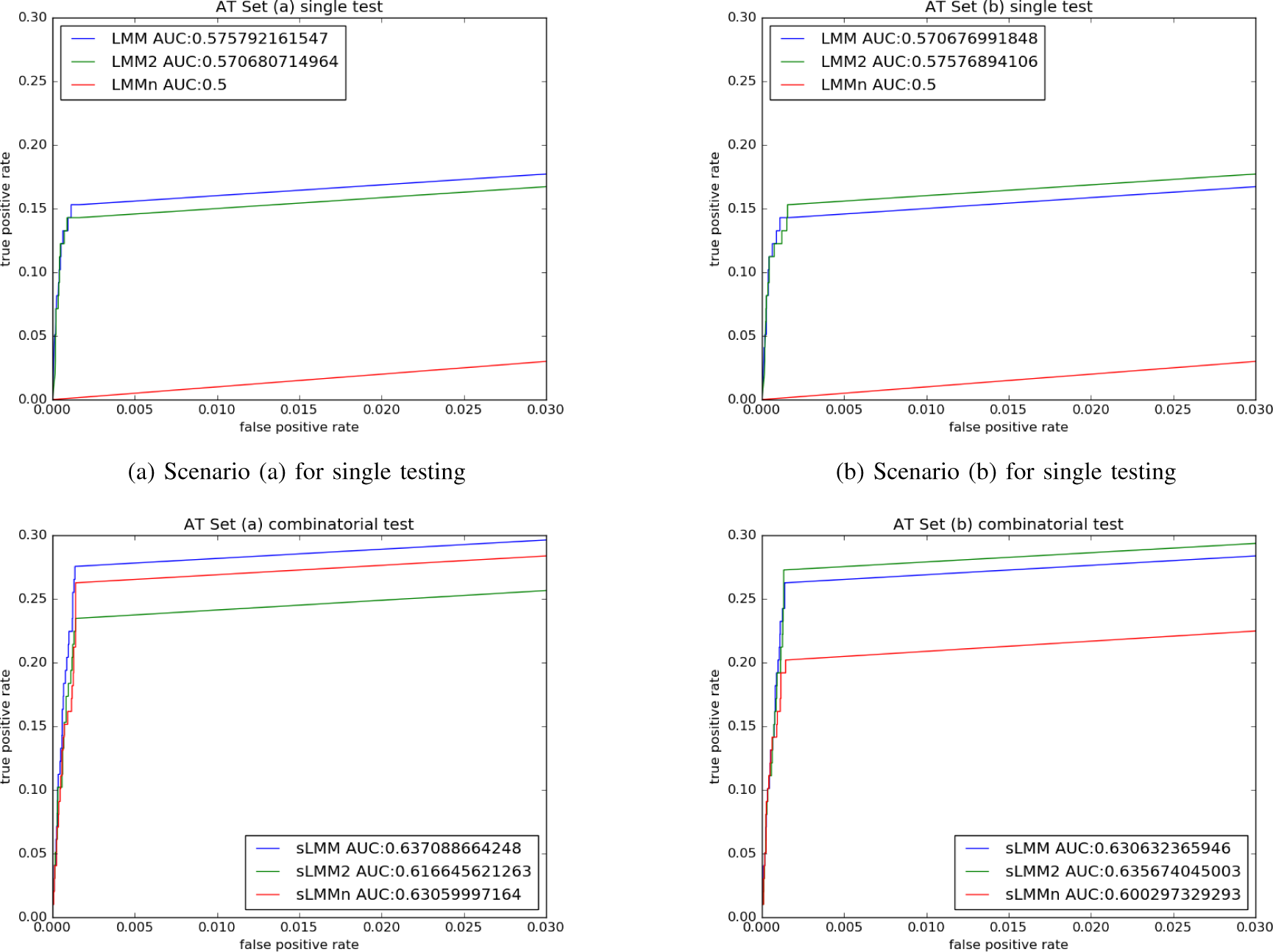
ROC curve to compare LMM^2^ with LMM (top row) and sLMM^2^ with sLMM (bottom row) on two different synthetic data sets

As the figure shows, for Case (a) where there is only a simple population structure, traditional LMM behaves better than our proposed higher order model. However, for Scenario (b), when we are simulating more complicated population structure which is more similar to the real dataset than Case (a), our proposed higher order LMM-LMM2 behaves best as the data is generated with second order kinship matrix in the figures we show.

### C. Semi-emperical Synthetic Data Set Experiment Two

The second experiment is an extension of the first one, and it is aimed to show that under different combinations of parameters governing the generation process of data, our proposed higher order LMM can consistently outperform previous methods.

#### 1) Data Generation

We generate synthetic data sets following the same way of Case (b) in the previous experiment, under Equation 6. Default values of parameters are set in the same way.

To make a fair comparison between these models, we repeat the data generation process with different parameters. In addition to these three parameters (*p_h_*, *p_s_*, *p_e_*), we also consider the number of causal SNPs sampled *N* (default value *N* = 100) as a parameter for data generation. Together with another parameter of the model *K* (default value *K* = 1000), which indicates the number of SNPs our model reports. We repeat our experiment many times with different combination of these five parameters.

The candidate values for each of these parameters are showed in Table II. Our experiment falls into five parts. In each part, we adjust one of those parameters while other parameters set to the default value. For each configuration of parameters, we repeat the experiment five times and report average area under ROC curve. We plot the curve of these averaged values along the different parameters of the parameter of interest.

**Table II:** Values of variables used for experiments

| parameter | Values |  |  |  |  |
| --- | --- | --- | --- | --- | --- |
| $p_e$ | 0.2 | 0.4 | 0.6 | 0.8 | 1.0 |
| $p_f$ | 0.2 | 0.4 | 0.6 | 0.8 | 1.0 |
| $p_s$ | 0.2 | 0.4 | 0.6 | 0.8 | 1.0 |
| $N$ | 100 | 500 | 800 | 1000 | 2000 |
| $K$ | 20 | 50 | 100 | 500 | 2000 |

#### 2) Results

Results are showed in Figure 5. As the figure shows, for single association mapping, LMM^2^ also shows some unstable behavior when its performance is compared to LMM, but in most cases (except experiment set of the parameter *p_f_*), LMM^2^ can outperform LMM. The figure also shows that LMM^*n*^ usually works better than LMM and LMM^2^ in single SNP testing cases. For combinatorial association mapping cases, sLMM^2^ outperform sLMM in almost all the cases. sLMM^*n*^ does not show a stable performance compared to sLMM, although its performance still exceeds sLMM in the majority of cases.

**Figure 5:**
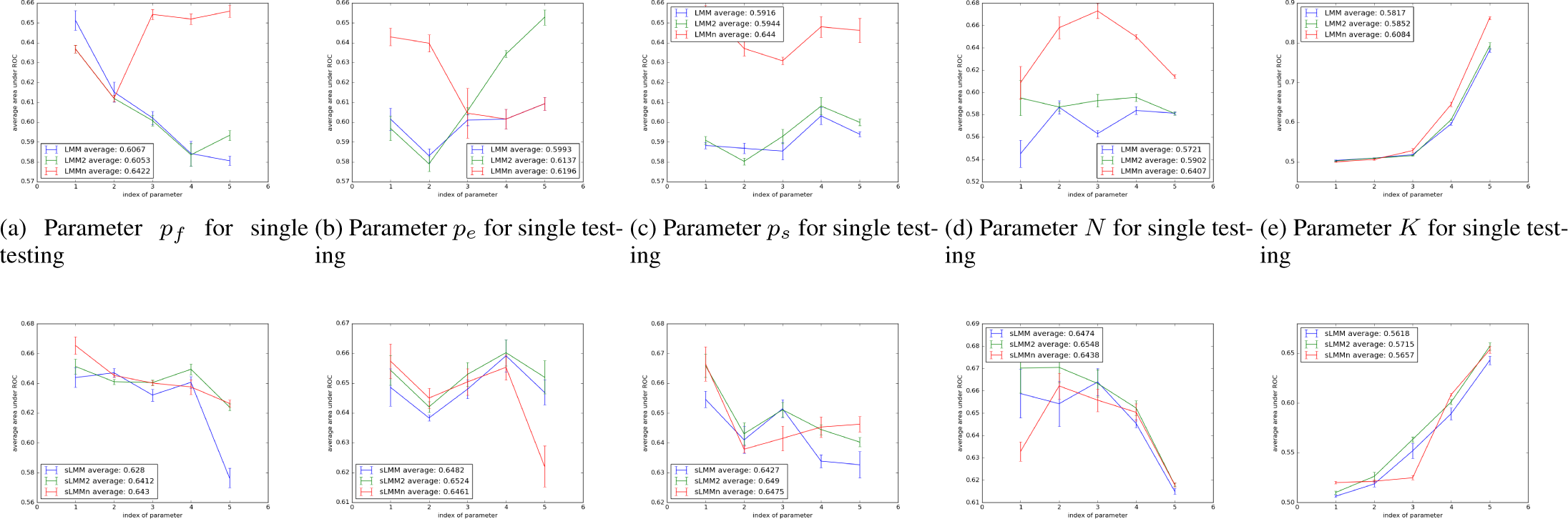
Curves to show the averaged area under ROC under different settings

It is also interesting to compare the differences between combinatorial association mapping models and single association mapping models. As the figure shows, combinatorial association mapping models (sLMM, sLMM^2^ and sLMM^*n*^) usually outperforms the corresponding single association mapping models (LMM, LMM^2^ and LMM^*n*^), which indicates the necessity of introducing combinatorial association mapping models to overcome the limitation of traditional hypothesis testing in GWAS tasks.

### D. Real Dataset Experiment

After showing the advantage of higher order LMM over synthetic data sets, we apply our proposed model real data set of *A.thaliana* where both genome information and phenotype information are available. To show the feasibility of our model compared to traditional LMM, we collect the candidate genes for each phenotype from [1].

Discarding the results that do not show a difference between original LMM and our proposed model, Table III shows the results of 10 sets of candidate genes and 20 phenotypes in correspondence. For the meaning of these phenotypes, please refer to [1].

**Table III:** Experiment Results on real data sets, each model is constrained to report the top *K* most significant associations discovered.

| Candidate Genes | Phenotype | Single Testing |  |  | Combinatorial Testing |  |  |
| --- | --- | --- | --- | --- | --- | --- | --- |
|  |  | LMM | LMM<sup>2</sup> | LMM<sup>n</sup> | sLMM | sLMM<sup>2</sup> | sLMM<sup>n</sup> |
| Anthocyanin | Anthocyanin 16 | 4 | **5** | 2 | 0 | 0 | 1 |
| Chlorosis | LY | 0 | **1** | 1 | 0 | **1** | 0 |
|  | Chlorosis 22 | 2 | 2 | 2 | 3 | **4** | 1 |
| Leaf-curl-roll | Leaf roll 10 | 1 | **2** | 1 | 2 | **3** | 0 |
| Dormancy | Secondary Dormancy | 0 | 0 | 0 | 0 | **1** | 0 |
| Flowering_time | LD | 3 | **4** | 3 | **4** | 2 | 3 |
|  | SD | **3** | 2 | 0 | 1 | 1 | **6** |
|  | FT10 | 4 | 5 | **8** | 7 | 7 | 5 |
|  | FT16 | 5 | 5 | 4 | 3 | 1 | **6** |
|  | FT22 | 2 | 2 | 2 | **6** | 5 | 4 |
|  | FLC | 4 | **5** | 4 | **6** | 5 | 5 |
| Pseudomonas-R-Genes | *avrRpt2* | 0 | 0 | 0 | 1 | **2** | 0 |
| Germination | Germ 16 | 0 | 0 | 0 | 2 | **4** | 0 |
|  | Germ 22 | 1 | 1 | 1 | 0 | **1** | 1 |
| Leaf_size | After Vern Growth | 1 | **2** | 1 | 2 | 2 | 0 |
| Senescence | FT Duration GH | 2 | 2 | 2 | **3** | 2 | 0 |
|  | LC Duration GH | 1 | 1 | 1 | 0 | 0 | **1** |
|  | LFS GH | 1 | 0 | 1 | 1 | 1 | 0 |
|  | RP GH | 1 | 1 | 1 | 0 | **2** | 0 |
| Silique | Silique 16 | 0 | **1** | 0 | 0 | **1** | 0 |
| Sum |  | 35 | 41 | 34 | 41 | 45 | 33 |

For Kinship matrix, we choose the group information to be the same as genome sequence, which leads to a kinship matrix of *XX^T^*.

Table III shows the true positives of the discoveries of each model, constraining each model to discover the same number of most significant associations. It shows that our proposed method, which raises the kinship matrix to the second order, behaves the best in both single SNP testing scenario and combinatorial testing scenario.

Interestingly, the results show that combinatorial testing scenario behaves better than its counterparts in a single testing scenario, which probably shows that most of these associations are a result of combinatorial effects contributed by a set of genetic markers, instead of one.

It is also noticeable that, for all these results reported, there are only five cases where LMM/sLMM behaves better than its squared counterpart. Out of these five cases, four of them are in the candidate gene set that is related to Flowering time. The reason we conjecture is that, whether the effect sizes of association should be explained in a more complicated way may depend on the phenotype of these associations of interest. Phenotypes regarding flowering time might be better explained with traditional 1st order LMM, for the reason that flowering time may depend more on the environment than genetics relationship and higher order kinship matrix may dilute the geographical information genetics captured.

#### 1) Results Comparison

Besides a numerical comparison of the discovered performance between these methods, here we select some typical examples to compare the genes each model discovered. The following associations discussed are verified with the online data base TAIR [26]^1^.

For example, for the phenotype that describes the visual chlorosis in plants under 22 °C after five weeks of growth (Row 3 in Table III), all three single testing methods discover the same two associated two genes: *AT4G24290* and *AT1G74710*, while for combinatorial testing, sLMM discovered one more associated gene *AT5G44030* and sLMM^2^ discovered one more in addition to what sLMM has discovered, namely *AT1G28380*. However, on the other hand, sLMM^*n*^ did not discover any of those associations, but it discovered *AT4G37000*.

For the phenotype that describes leaf roll presence 8 weeks post germination for plants grown at 10 °C (Row 4 in Table III), LMM and LMM^*n*^ discovered the association *AT1G09530* and LMM^2^ discovered one more associated gene (*AT3G14110*) additionally. On the other hand, combinatorial association methods discovered completely different sets of genes, sLMM found *AT3G56400* and *AT3G02570* and sLMM^2^ discovered *AT1G34210* additionally.

For the phenotype that describes the days from removal from stratification (3 days at 4 °C in the dark) until emergence of first cotyledon at 16^0^C (Row 13 in Table III), single testing methods did not discovered any associated genes. sLMM discovered two associated genes, namely *AT2G29630* and *AT3G54810* and sLMM^2^ discovered two additional associated genes, namely *AT5G51760* and *AT2G18790*.

These examples all validate the performance of our proposed squared LMM that, by raising the order of kinship matrix, the model can typically discover more associated genes in addition to what original LMM can discover. In fact, except the phenotypes that are related to flower timing and few other cases, squared models are always capable of discovering associated genes in addition what original models discover in both single testing case and combinatorial case.

## IV. Conclusion

In this paper, we proposed an extension of linear mixed model to consider more complex confounding factor structures and this improvement can be applied to both traditional linear mixed model that tests the association individually and sparse linear mixed model that test the combinatorial association.

We first made the argument that raising the order of kinship matrix can gain many advantages of the model including 1) allowing the model to consider hidden confounding structures 2) distinguish the statistical representation power for fixed effect variables and kinship matrix when kinship matrix is calculated as the covariance matrix of fixed effect variables 3) can be reformulated into another model that considers that the random effects are not independently, but also following a covariance matrix. Based on these reasonings, we proposed our model LMM^2^ for single association testing case and sLMM^2^ for combinatorial association testing case and further extend these models to LMM^*n*^ and sLMM^*n*^ for comparison.

As our extensive experiment on synthetic data showed, our squared model can outperform its counterparts for correctly discovering the causal SNPs on both simulations and real data in most cases. However, for flowering time-related phenotypes in real data, our proposed model discovered less associated genes than the traditional linear mixed model.

Since this model is primarily an improvement over previous models in the performance of confounding correction. In the future, we plan to proceed to incorporate more powerful regularizers for multivariate regression to further improve the performance, like the Precision Lasso [9], which is used to account for linkage disequilibrium of SNPs. To further improve the modeling performance of associations, we will also consider the deep neural network approaches [27]. A neural network equivalent of vanilla linear mixed model has already been proposed [28], but more efforts are necessary to incorporate our higher order linear mixed model. We will also implement our method into the GWAS tool box GenAMap [29]^2^ for convenient usage of biologists.

## Acknowledgement

This material is based upon work funded and supported by the Department of Defense under Contract No. FA8721-05-C-0003 with Carnegie Mellon University for the operation of the Software Engineering Institute, a federally funded research and development center. This work is also supported by the National Institutes of Health grants R01-GM093156 and P30-DA035778.

## Footnotes

1 http://www.arabidopsis.org/

2 http://genamap.org/

